# Mapping Chromatin Occupancy of *Ppp1r1b-lncRNA* Genome-Wide Using Chromatin Isolation by RNA Purification (ChIRP)-seq

**DOI:** 10.1101/2023.11.04.565657

**Authors:** John Hwang, Xuedong Kang, Charlotte Wolf, Marlin Touma

## Abstract

Long non-coding RNA (lncRNA) mediated transcriptional regulation is increasingly recognized as an important gene regulatory mechanism during development and disease. LncRNAs are emerging as critical regulators of chromatin state; yet the nature and the extent of their interactions with chromatin remain to be fully revealed. We have previously identified *Ppp1r1b-lncRNA* as an essential epigenetic regulator of myogenic differentiation in cardiac and skeletal myocytes in mice and humans. We further demonstrated that *Ppp1r1b-lncRNA* function is mediated by the interaction with the chromatin-modifying complex polycomb repressive complex 2 (PRC2) at the promoter of myogenic differentiation transcription factors, *TBX5* and *MyoD1*. Herein, we employed an unbiased chromatin isolation by RNA purification (ChIRP) and high throughput sequencing to map the repertoire of *Ppp1r1b-lncRNA* chromatin occupancy genome-wide in the mouse muscle myoblast cell line. We uncovered a total of 99732 true peaks corresponding to *Ppp1r1b-lncRNA* binding sites at high confidence (*P*-value < 1e-5 and enrichment score ≥ 10). The *Ppp1r1b-lncRNA*-binding sites averaged 558 bp in length and were distributed widely within the coding and non-coding regions of the genome. Approximately 46% of these true peaks were mapped to gene elements, of which 1180 were mapped to experimentally validated promoter sequences. Importantly, the promoter-mapped binding sites were enriched in myogenic transcription factors and heart development while exhibiting focal interactions with known motifs of proximal promoters and transcription initiation by RNA polII, including TATA, transcription initiator, CCAAT-box, and GC-box, supporting *Ppp1r1b-lncRNA* role in transcription initiation of myogenic regulators. Remarkably, nearly 40% of *Ppp1r1b-lncRNA*-binding sites mapped to gene introns, were enriched with the Homeobox family of transcription factors, and exhibited TA-rich motif sequences, suggesting potential motif specific *Ppp1r1b-lncRNA*-bound introns. Lastly, more than 136521enhancer sequences were detected in *Ppp1r1b-lncRNA*-occupancy sites at high confidence. Among these enhancers,12% exhibited cell type/tissue-specific enrichment in fetal heart and muscles. Together, our findings provide further insights into the genome-wide *Ppp1r1b-lncRNA:* Chromatin interactome that may potentially dictate its function in myogenic differentiation and potentially other cellular and biological processes.

## INTRODUCTION

The majority of the mammalian genome is transcribed to produce RNA transcripts, most of which display no protein-coding potential [**1**]. Long non-coding RNA (lncRNA) transcripts define an expanding class of non-coding RNA species that are longer than 200 nucleotides and lack functional open reading frames. Like mRNAs, lncRNAs are primarily transcribed by RNA polymerase II (RNA Pol-II), 5′-capped, poly A-tailed, and post-transcriptionally modified mostly by splicing [**2,3**].

LncRNAs are pervasively transcribed across the genome and have emerged as important transcriptional regulators, affecting all layers of transcriptome regulation, including RNA transcription, splicing, and metabolism [**2–5**]. As our understanding of biochemical properties and functional diversity of lncRNA continues to evolve, it is widely accepted that lncRNAs can exert diverse functions that arise from their ability to form complex secondary structures with DNA-, RNA-, and protein-binding abilities, leading to complex RNA-DNA, RNA-RNA, or RNA-protein interactions [**5,6**]. Moreover, a single lncRNA can contain several binding loops that are able to bind to nucleic acids via base pairing or to proteins by certain RNA binding motifs, thus allowing the coordination of signals between different types of macromolecules and chromatin-modifying complexes [**5,6**]. It has been evident that several lncRNAs, such as HOTAIR and Braveheart, can execute their regulatory functions by recruiting chromatin modification complexes and altering the state of chromatin accessibility, leading to transcriptional activation or repression [**7, 8**]. By performing these diverse functions, lncRNAs can influence cellular biology, molecular processes, and tissue homeostasis at multiple levels, including transcriptome regulation, molecular networking, cellular differentiation, and developmental decisions [**2–8**].

During development, chromatin states are the key determinants of cellular differentiation, identity, and fate [**9–12**]. We have previously identified *Ppp1r1b-lncRNA* as an essential and functionally conserved epigenetic regulator of myogenic differentiation of cardiac and skeletal myocytes in both mice and humans [**13**]. Importantly, in response to *Ppp1r1b-lncRNA* loss, human induced pluripotent stem cells (hiPSCs)-derived cardiac progenitors and skeletal myoblast cell lines failed to produce early markers of myogenic differentiation program upon induction [**13**]. Cellular differentiation requires activation of specific transcriptional programs that are governed by cell-specific master regulators and transcription factors [**14,15**]. We have demonstrated that *Ppp1r1b-lncRNA* interferes with polycomb repressive complex 2 (PRC2) binding at target promoters of the master transcription factors of myogenic differentiation, *TBX5* and *MyoD1*, leading to decreased enrichment of H3K27me3, a PRC2-catalyzed epigenetic marker of transcriptional repression. In turn, the resulting enhanced chromatin accessibility leads to positive regulation of *TBX5* and *MyoD1* and induction of myogenic differentiation programs in cardiac and skeletal myocytes. These findings support the key role of *Ppp1r1b-lncRNAs* in modulating chromatin states in a gene-specific manner to promote myogenic differentiation.

Interestingly, while *Ppp1r1b-lncRNA* was initially thought to act locally on a neighboring protein-coding gene [**3**], our mechanistic studies, including chromatin isolation by RNA purification-polymerase chain reactions (ChIRP-PCR), revealed that *Ppp1r1b-lncRNA* executes its function by physically interacting with distantly located transcription factors (*TBX5* and *MyoD1).* In our work presented here, we set out to elucidate the full panel of *Ppp1r1b-lncRNA*-binding sites and explore how the specificity for *Ppp1r1b-lncRNA* interactions is achieved [**13**]. We employed a ChIRP strategy followed by single-read high throughput DNA sequencing and subsequent bioinformatics tools to map *Ppp1r1b-lncRNA* occupancy at the genome scale. By applying downstream peak calling pipeline and peak mapping to gene elements, we revealed genome-wide Ppp1r1b-lncRNA-bound chromatin and gained further insights into the specific motifs that may underlie *Ppp1r1b-lncRNA* function at proximal promoters or distant enhancers of its putative target genes, including those encoding myogenic differentiation factors, transcription regulation, and chromatin modifiers.

## METHODS

### ChIRP Assay

#### I. Probe design for ChIRP

The Magna ChIRP RNA interactome kit (EMD Millipore Corp.) was used. Assays were performed per manufacturer’s protocol. The capture probe is an antisense-oligo high-affinity probe, targeted against a unique *Ppp1r1b-lncRNA* sequence, was designed using http://www.singlemoleculefish.com. The probe was compared with the mouse genome using the BLAT tool and no noticeable homology to non-*Ppp1r1b-lncRNA* targets were detected. An anti-sense oligo probe against lacZ RNA was provided by the ChIRP kit and used as a negative control. Both probes were biotinylated at the 3’ end.

#### II. Cell culture

Mouse myoblasts, C2C12 cell line, were cultured in DMEM (Invitrogen) supplemented with 10% Fetal Bovine Serum (FBS) and 1% Pen/Strep (Invitrogen).

#### III. Cross-linking, sonication, and hybridization

C2C12 cells were grown to log-phase in tissue culture plates and rinsed once with room temperature Phosphate buffer saline (Pbs). Cells were treated with glutaraldehyde for cross-linking as described previously [**13, 16, 17**]. The resulting chromatin was fragmented by sonication. A sample consisting of 2% of the total input chromatin was then removed and served as the sequencing control. A biotinylated complementary anti-sense oligo probe was hybridized to *Ppp1r1b-lncRNA* and then isolated using magnetic streptavidin beads. No cross-hybridization with the LacZ probe was detected. The co-purified *Ppp1r1b-lncRNA* bound chromatin was then eluted for protein, RNA, and DNA. Using a combination of RNase A and RNase H, the DNA was gently eluted off of beads as described by the manufacturer’s instructions and processed into small fragments for library preparation.

### ChIRP-Seq

#### I. Library preparation and high-throughput sequencing

The sequencing libraries were constructed from the ChIRP-captured and control “input” DNA fragments. Around 3 ng *Ppp1r1b-lncRNA*-ChIRP DNA and 3 ng control DNA “input” were used for library preparation as per manufacturers’ protocol. DNA fragments were subjected to DNA-end repair, 3’-ad overhanging, and adaptors ligation, and then amplified using PCR. After size selection (between 100 and 500 bp), qualified *Ppp1r1b-ncRNA*-ChIRP and control DNA libraries were used for high throughput single-end (SE) sequencing on BGIseq at a read length of 50 bp, generating an average of 38 million raw sequencing reads per sample.

#### II. Bioinformatic analysis workflow

##### 1. Data filtering

Raw sequencing data were filtered using the software Short Oligonucleotide Analysis Package (SOAP) nuke to remove adapter sequences, contamination, and low-quality reads. The following parameters were used for the SOAPnuke filter: -l 5 -q 0.5 -n 0.1 -Q 2 -c 40. Reads were considered “low-quality” if any of the following was true: 1) the ratio of N (unmappable reads) in whole read was >10%; 2) reads in which unknown bases exceeded 10%; or 3) the ratio of base whose quality was less than 20 was > 10%.

##### 2. Reads alignment

Clean reads that passed quality check measures were stored in FASTQ format and then aligned to the reference genome GRCm38/mm10 (Genome Reference Consortium Mouse Build 38 Organism: Mus musculus 10) using SOAP aligner SOAP2 (Version: 2.21t) [**18**]. No more than two mismatches were allowed in read alignment. Base coverage was normalized per million mappable reads. Reads from *Ppp1r1b-lncRNA*-ChIRP and control samples were aligned separately. The alignment results were then used for peak calling.

##### 3. Peak calling and identifying true peaks

The uniquely mapped clean reads resulting from the alignment step were then used for peak calling. Candidate peaks for each sample were called using the software Model-Based Analysis for ChIP-Seq (MACS) v1.4.2. [**19**]. The following parameters were used for peak calling: -g mm

-s 50 -p 1e-5 -m 10 30 --broad -B –trackline:

-g: mappable genome size, defined as the genome size that can be sequenced. GRCm38/mm10= 1.87e9 (G)

-s: size of sequencing tags

-p: *p*-value cutoff: 1e-5

-m: minimum length of called peak (10) and maximum gap allowed between two peaks (30) to be merged. 14: smallest peak size

--: input file format in BED format

Based on λlocal, MACS workflow uses dynamic Poisson distribution to calculate the *p*-value of the specific region based on the unique mapped reads. The region is defined as a peak when the *p*-value < 1e-5 (by default). The MACS-predicted peaks are also assigned enrichment scores. The more enriched these regions are, the more likely they represent true binding sites. In our analysis, MACS-predicted peaks were further filtered to obtain a list of true peaks (true *Ppp1r1b-lncRNA*-binding sites) at stringent enrichment score values of >=10.

##### 5. Peak mapping to promoter elements

To identify peaks that overlap with promoters, 25111 coding and 3077 non-coding promoters for GRCm38/mm10 were downloaded from the Eukaryotic Promoter Database new (EPDnew) (https://epd.epfl.ch/epdnew/documents/MmNC_epdnew_001_pipeline.php) [**20, 21**]. The EPDnew promoters are experimentally validated with next-generation sequencing-based whole-genome TSS mapping protocols, such as Cap Analysis of Gene Expression (CAGE) and Oligocapping, and include TATA box, Initiator motif, CCAT, and other well-established promoter elements. Using the Bedtools “*intersect*” feature, true peaks (enrichment score ≥ 10) with at least 30% overlap with EPDnew promoters were identified.

##### 7. Peaks mapping to putative enhancer elements

Putative enhancer elements were obtained from EnhancerAtlas Browser (EnhancerAtlas 2.0; http://www.enhanceratlas.org/indexv2.php) [**22**]. The database provides enhancer annotation in nine species, including human, mouse, fly, worm, zebrafish, rat, yeast, chicken, and boar annotations. The consensus enhancers were predicted based on multiple high throughput experimental datasets (e.g. histone modification, CAGE, GRO-seq, transcription factor binding, and DHS). Currently, the updated database contains 6,198,364 enhancers and 7,437,255 enhancer-gene interactions involving 31,375 genes for 241 murine tissue/cell types identified from 5,838 datasets such as NCBI GEO datasets, ENCODE project portal at UCSC, Epigenome Roadmap and FANTOM5. As above, we used the Bedtools “*intersect”* feature to identify true peaks that overlapped with the enhancer elements at confidence score >=1

##### 8. Peak visualization using UCSC Genome Browser and Broad Institute Integrative Genomics Viewer (IGV)

The UCSC Genome Browser, which contains genome references assemblies for multiple species, was used to visualize and download *Ppp1r1b-lncRNA-ChIRP* derived peaks as well as specific genes and regions genome-wide. After selecting the GRCm38/mm10 genome on the UCSC genome browser, *Ppp1r1b-lncRNA-ChIRP,* control, and Peak Bed files were uploaded to custom tracks. The distribution of peaks across the genome and within specific regions was shown. IGV was used in a similar fashion for visualization and analysis [**23**].

##### 9. Motif analysis using MEME-SEA

Motifs analysis was performed using Multiple EM for Motif Elicitation (MEME)-Simple Enrichment Analysis (SEA) v5.5.3. [**24,25**], using the following command line: sea --verbosity 5 --oc --thresh --align center --p *input_file* --m *motif_database* 10.0. MEME works by searching for repeated, ungapped sequence patterns that occur in the DNA or protein sequences provided by the user. The motifs discovered motifs can be compared with databases of known motifs, identify matches to the motifs, and display the motifs in various formats. The motif database used in this study is the Universal PBM Resource for Oligonucleotide Binding Evaluation (UniPROBE) database for the murine species, which is generated by universal protein binding microarray (PBM) assays on the *in vitro* DNA binding specificities of proteins [**26**].

#### III. Functional enrichment of peaks’ related genes

##### 1. Identifying peaks’ related genes

Data was downloaded from the UCSC Genome browser. To identify peaks’ related genes, we applied the following criteria: A. The reads are uniquely mapped to a protein-coding gene. B. The genes must be annotated (gene name present). C. The gene status is known.

##### 2. Gene ontology annotation of peaks’ related genes

Gene Ontology (GO) analysis was used to predict the main biological functions that are enriched in the peaks’ related genes and assign them to specific molecular functions, biological processes, and cellular components [**27**]. All peaks’ related genes were mapped to GO terms in the database (http://www.geneontology.org). The number of genes for every term was then calculated. Finally, a hypergeometric test was used to find significantly enriched GO terms in the query list of peaks’ related genes. The calculated *p*-value goes through Bonferroni correction and a corrected p-value ≤0.05 defines the significantly enriched GO terms in the peaks’ related genes.

##### 3. KEGG pathway enrichment

To further understand the biological functions of the peak-related genes in a pathway-based contest, KEGG (Kyoto Encyclopedia of Genes and Genomes) was used to perform pathway enrichment analysis [**28**]. This analysis identifies significantly enriched metabolic pathways or signal transduction pathways in peak-related genes compared with the target regions’ background. The analysis process follows the same pipeline as that in GO analysis.

### Statistical Analysis

Quantified results and statistical parameters for each bioinformatic analysis step were presented with their data within the corresponding sections of the text.

## RESULTS

### I. Quality Control and Alignment Statistics Results

In this study, we mapped *Ppp1r1b-lncRNA* chromatin occupancy genome-wide by ChIRP-seq in mouse myoblast cell line, which expresses endogenous *Ppp1r1b-lncRNA* [**3, 13**]. Sequencing libraries were prepared from *Ppp1r1b-lncRNA*-ChIRP and control input DNA fragments and subjected to single read high throughput sequencing at a read length of 50 bp [**Figure 1**]. An average of 38 million raw sequencing reads per sample were generated. After filtering low-quality reads and removing adaptor sequences, 36534935 and 38429369 (99.13% and 98.04%) clean reads were obtained from *Ppp1r1b-lncRNA-ChIRP* and control samples, respectively, to be used for downstream analysis. The sequencing data summary for each sample is summarized in [**Table 1**].

**Figure 1.**
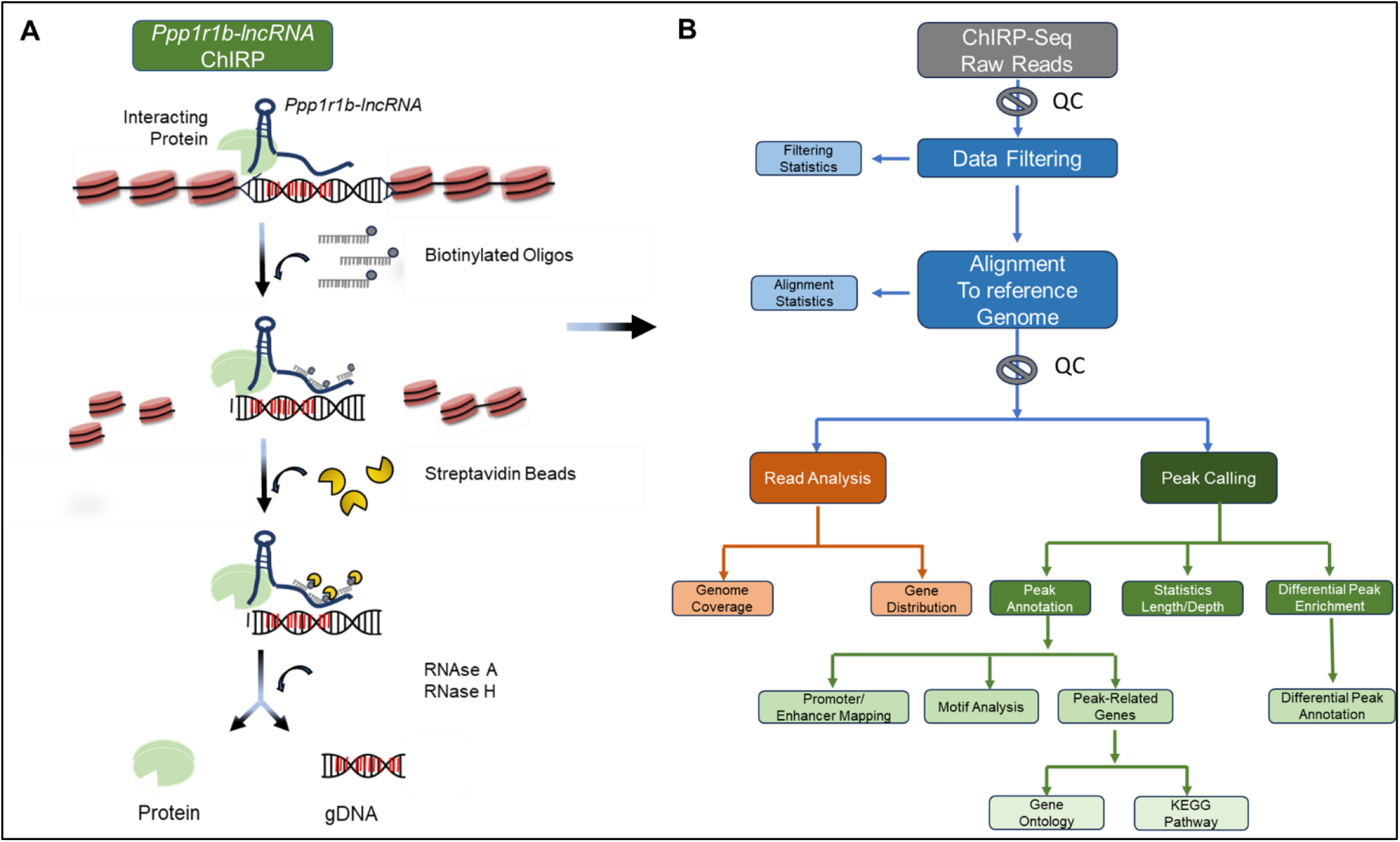
Schematic Summary of *Ppp1r1b-lncRNA*-ChIRP Assay and Bioinformatic Pipeline. **A.** Chromatin was crosslinked *in vivo.* Biotinylated tiling probes were hybridized to target lncRNA, and chromatin complexes were purified using magnetic streptavidin beads, followed by stringent washes. The lncRNA-bound DNA or proteins were eluted with a cocktail of RNase A and H. A putative lncRNA binding DNA sequence is highlighted in red. Adopted from Chu *et al.* 2011. **B.** The ChIRP-captured DNA fragments and control input libraries were subjected to next-generation sequencing. After quality control (QC) and alignment to the reference genome, the clean uniquely mapped reads were used for peak calling and downstream bioinformatic analysis.

**Table 1 (Excel Sheet 1):** Summary of sequencing data for each sample.

| Sample ID | Fragment Length(bp) | Sequencing Strategy | Clean Reads Number | Clean Data Size (bp) | Clean Rate(%) |
| --- | --- | --- | --- | --- | --- |
| *Ppp1r1b-lncRNA*_ChIRP | 100-500 | SE50 | 36534935 | 1.83E+09 | 99.13 |
| Control | 100-500 | SE50 | 38429369 | 1.92E+09 | 98.04 |

Footnotes: Fragment Length (bp): DNA fragment for library building; SE50: Strategy to sequencing sample with single end (SE), and the following number reflects read length. Clean Reads Number: The count of reads number in clean data. Clean Data Size: The count of bases in clean data. Clean Rate (%): The ratio of clean data size to raw data size=Clean Data Size (bp)/Raw Data Size (bp).

For read mapping, the clean reads were mapped to the mouse reference genome GRCm38/mm10 using SOAPaligner/soap2 [**18**]. Only the alignments within 2 bp mismatches were considered and strict quality control measures for each sample were applied achieving a genome mapping rate of 95.21% and 96.97% for *Ppp1r1b-lncRNA-ChIRP* and control samples, respectively [**Supplemental Table 1**].

For peak calling, only the uniquely mapped clean reads that only map to one genomic position in the total reads were included. Following this criteria, 31664010 and 32579793 uniquely mapped reads (86.7% and 84.78%) were obtained from *Ppp1r1b-lncRNA ChIRP* and control samples, respectively to be used for downstream peak calling and subsequent analysis. Alignment statistics results and genome mapping rate for each sample are summarized in [**Table 2**].

**Table 2 (Excel Sheet 2):** Alignment statistics results and genome mapping rate for each sample.

| Sample ID | Species | Clean Reads | Mapped Reads | Mapped Rate(%) | Uniquely Mapped Reads | Uniquely Mapped Rate(%) |
| --- | --- | --- | --- | --- | --- | --- |
| *Ppp1r1b-lncRNA*-ChIRP | mm10 | 36534935 | 34783233 | 95.21 | 31674010 | 86.7 |
| Control | mm10 | 38429369 | 37264694 | 96.97 | 32579793 | 84.78 |

Footnotes: Clean Reads: Total clean reads number; Mapped Reads: Total reads that can be mapped to the reference genome. Mapped Reads Rate (10%): The proportion of reads that can be mapped to the reference genome in total reads. Uniquely Mapped Reads: Total reads that only map to one position in the reference genome. Uniquely Mapped Rate (%): The proportion of reads that only map to one position in total reads.

### II. Genome and Gene Depth Distribution Analysis

The uniquely mapped reads that passed quality control measures were then used to estimate the genome depth distribution for each sample separately using BEDTools. The percentage of genome coverage for each sample is shown in [**Figure 2. A and B**]. Gene depth distribution was also obtained for each sample separately by BEDTools, and only those uniquely mapped reads were used in this analysis. As shown in [**Figure 2. C and D**], the average depth of *Ppp1r1b-lncRNA-ChIRP* reads exhibited differential distribution in relation to genic regions with increased coverage around TSS and towards the distal 50% part of genes.

**Figure 2.**
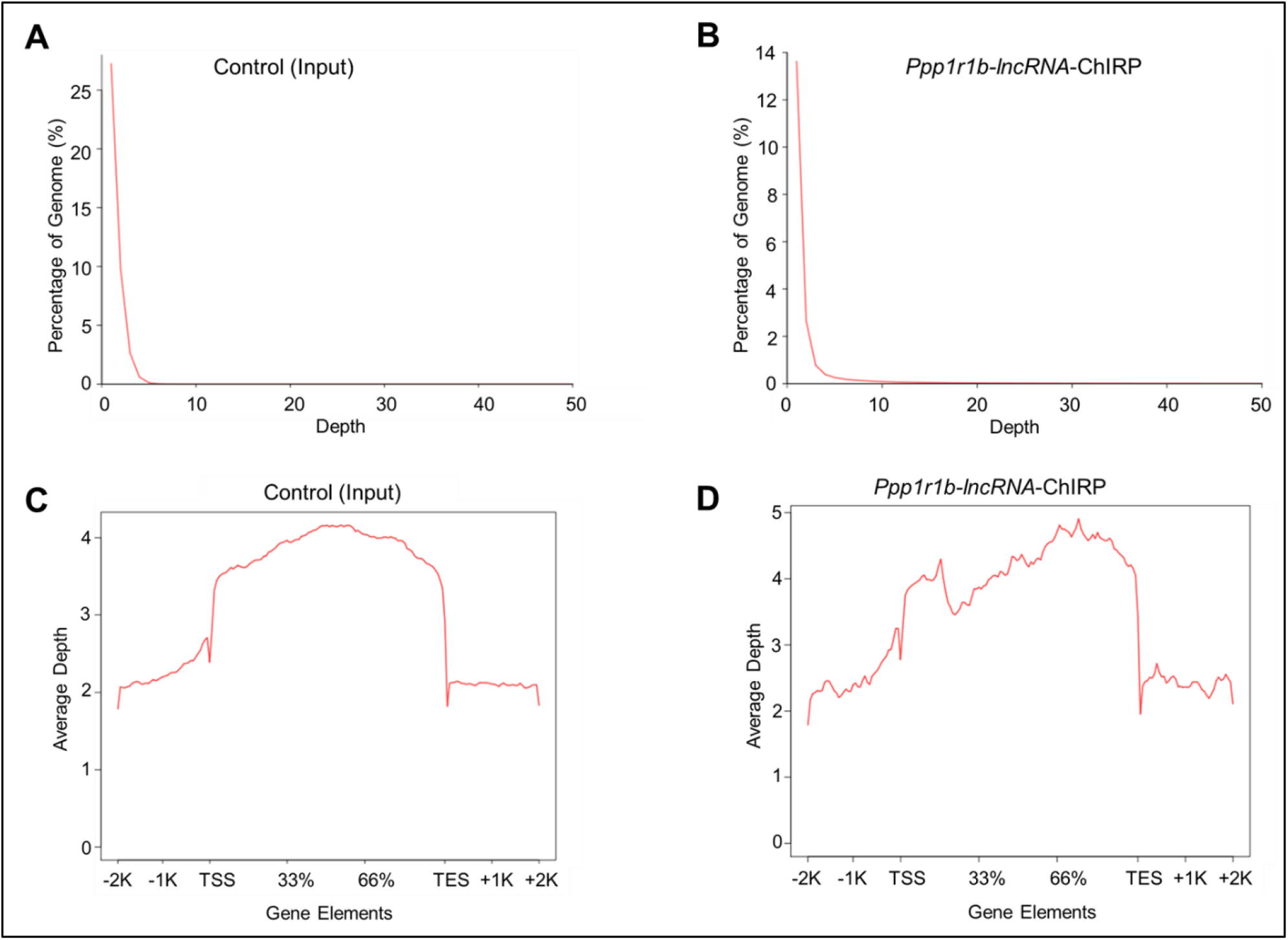
Genome and Gene Depth Distribution for Each Sample. **A and B.** Composite genome coverage of *Ppp1r1b-lncRNA*-ChIRP and Control (input) sequencing reads in mouse myoblasts cell line. These figures represent all reads in each sample. **C and D.** Composite sequencing reads profiles of *Ppp1r1b-lncRNA*-ChIRP and Control (input) samples across gene regions in mouse myoblasts cell line. These figures only represent the reads mapped to gene elements. The ‘Input’ library was used as a control.

### III. Peak Calling

The uniquely mapped reads were then used for genome-wide peak calling using MACS standard pipeline [**19**]. MACS-detected peaks statistics at a *p-*value cut-off <1e-5 are summarized in [**Table 3**]. MACS peak calling statistics results include peak location, peak enrichment score, and peak length.

**Table 3 (Excel Sheet 3):** Peak calling statistics.

| Sample ID | Peak Number | Total Length | Average Length | Total Tag Depth | Average Tag Depth | Genome Rate(%) |
| --- | --- | --- | --- | --- | --- | --- |
| *Ppp1r1b-lncRNA*-ChIRP | 261455 | 146128139 | 558 | 3848968.705 | 14 | 5.35 |

Footnotes: Peak number: Number of all MACS-detected peaks. Total Length: Total length of all peaks. Average Length: Average length of all peaks. Total Tag Depth: Total tag depth of all summits. Average Tag Depth: Average tag depth of all summits. Genome Rate: Proportion of total length of all peaks in the whole genome.

In total, MACS identified 261455 peaks in the *Ppp1r1b-lncRNA*-ChIRP sample that passed a *P* value < 1e-5 against control input. *Ppp1r1b-lncRNA-ChIRP* peaks were short and focal, ranging between 165-4500 bp in length with a mode of 165 bp and an average peak length of 558 bp [**Figure 3. A and B**]. With a genome coverage rate of 5.35%, the peaks were widely distributed in the intergenic-(53.4% of all peaks) and the genic-(46.4% of all peaks) regions. With reference to the gene elements, 39.6% of the gene-mapped peaks were located in introns, while 2.5% were mapped to exons, 2.2% mapped to immediate Up2k, and 2.1% mapped to immediate Down2K, of the TSS and TES, respectively [**Figure 3. C**].

**Figure 3.**
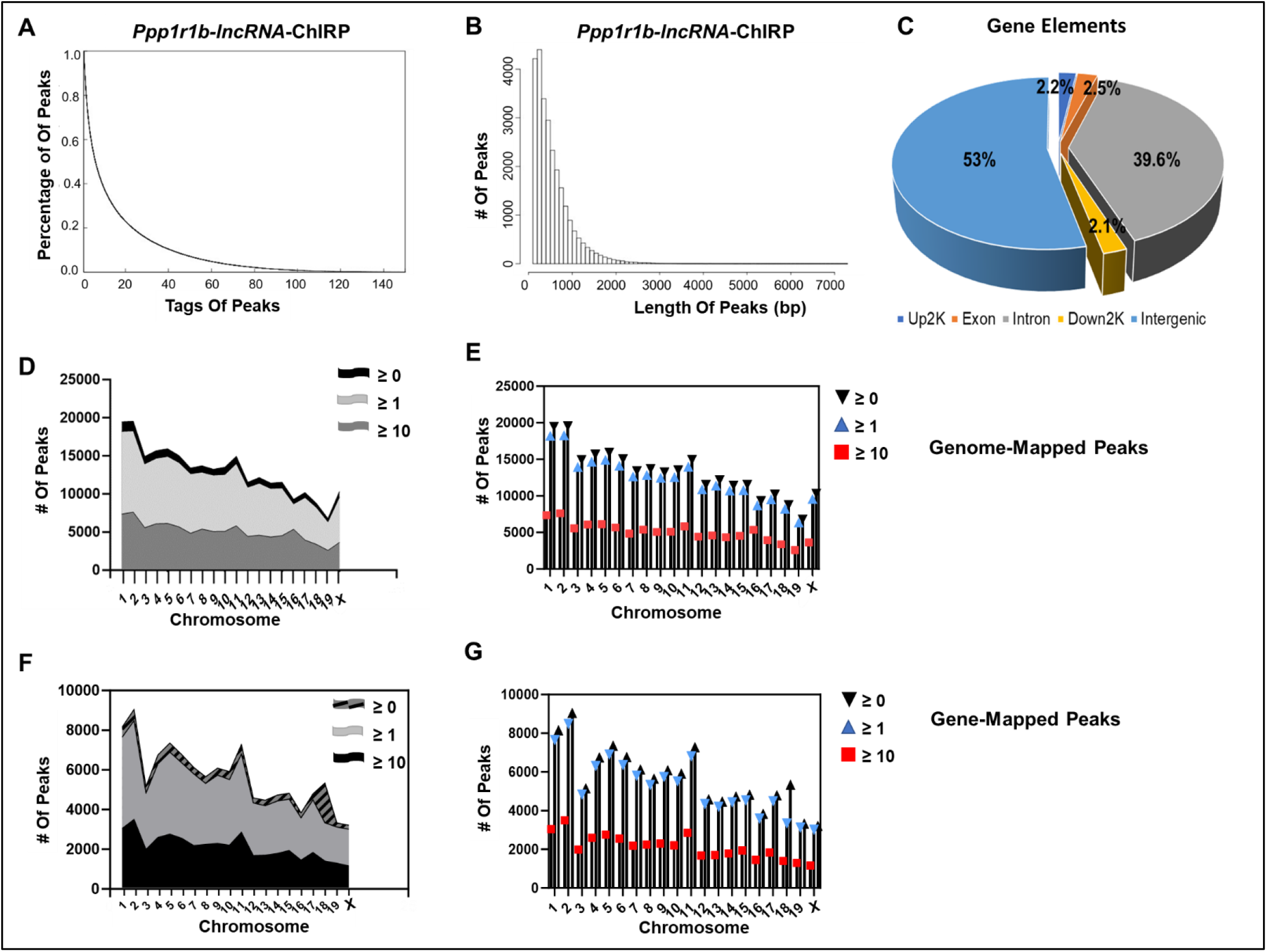
MACS-derived *Ppp1r1b-lncRNA-*ChIRP Peaks Statistics. **A.** Depth Distribution of all peaks called by MACS at *P*-value<1e-5. The X-axis indicates the number of reads, and the Y-axis indicates the proportion of peaks in the specific number of reads. **B.** Length distribution of all MACS*-*called peaks. The X-axis refers to peak length, and the Y-axis refers to peak numbers. **C.** Distribution of all MACS-called peaks based on genomic position and gene elements: intergenic, introns, downstream, upstream, and exons. **D and E.** Number of MACS-called Peaks (P<1e-5) genome-wide (**D**) and on each chromosome (**E**) at different enrichment score cut-off values (≥0, ≥1, and ≥10). The X-axis indicates chromosome number, and the Y-axis indicates the number of peaks mapped to each chromosome. **F and G.** Number of MACS-called peaks (P<1e-5) that mapped to known protein-coding genes on all chromosomes (**F**) and on each chromosome (**G**) at different enrichment score cut-off values (≥0, ≥1, and ≥10). The X-axis indicates chromosome number, and the Y-axis indicates the number of gene-mapped peaks on each chromosome.

To enhance the specificity of peaks that represent true *Ppp1r1b-lncRNA* binding sites, only peaks with an enrichment score equal to or greater than 10 were defined as true peaks that represent *Ppp1r1b-lncRNA*-binding sites and retained for further analysis. Using these parameters 99732 true *Ppp1r1b-lncRNA*-binding sites were identified genome-wide, of which 42393 (43% of true peaks) were mapped to protein-coding genes, indicating that the ratio of the gene-mapped peaks to the total number of peaks did not change significantly despite using more stringent thresholds for true peaks selection. Notably, like the genome-mapped binding sites [**Figure 3. D and E**], the gene-mapped binding sites [**Figure 3. F and G**] were widely spread on all chromosomes and retained similar patterns of distribution at different cut-off values for enrichment scores. Furthermore, the length distributions as well as enrichment score proportions of the genome-mapped binding sites [**Figure 4. A-D**] were comparable to those mapped to the genes [**Figure 4. E-H**]. The peak calling results were stored in wiggle files and viewed on the UCSC genome browser and the Integrative Genome Viewer (IGV) for peak visualization. Examples of *Pppp1r1b-lncRNA*-binding sites are presented in [**Supplemental Figure 1**].

**Figure 4.**
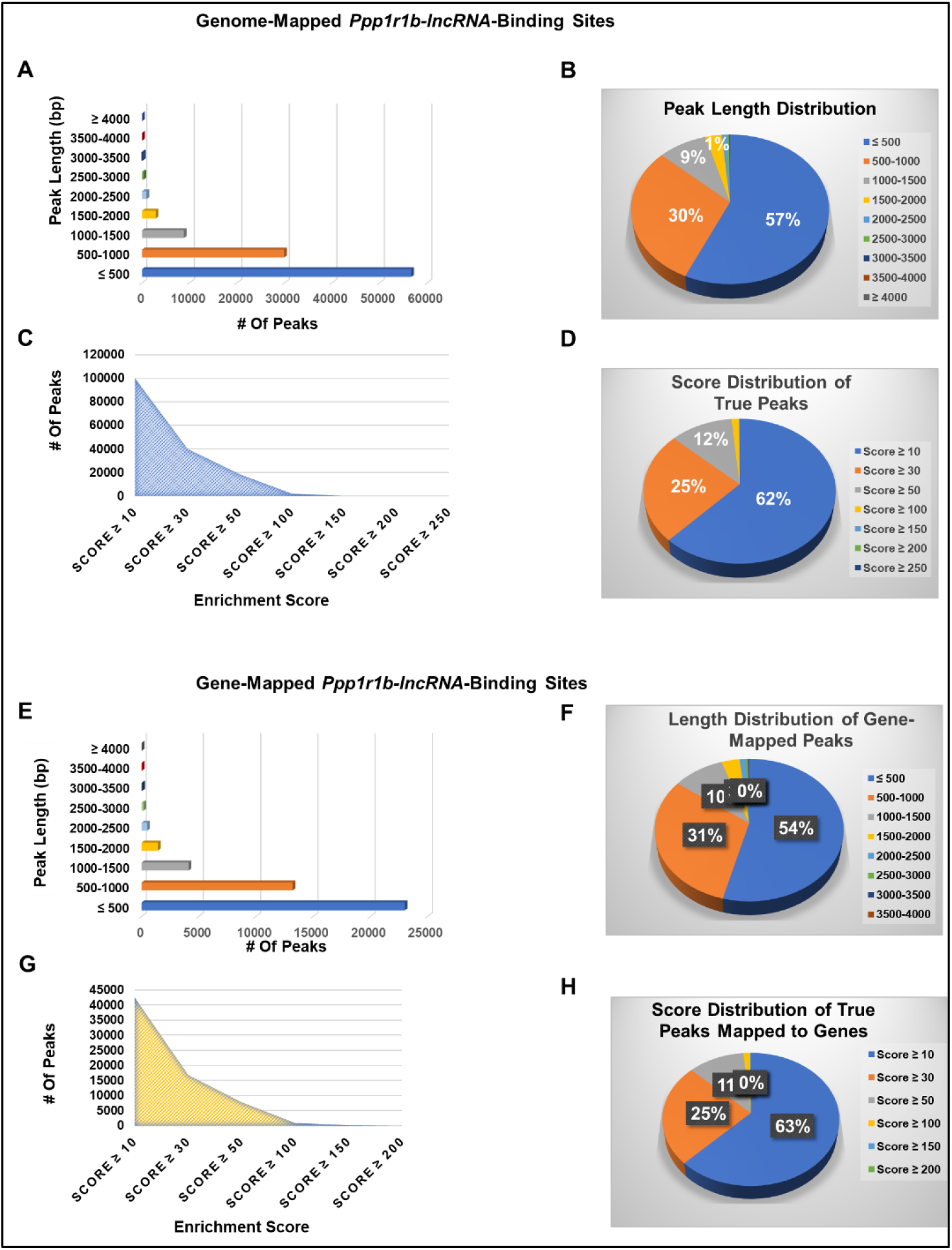
*Ppp1r1b-lncRNA*-Binding Sites (True Peaks) Statistics. **A.** Number of genome-mapped true peaks at different peak length (bp) values. **B.** Distribution of genome-mapped true peaks at different peak length (bp) values. **C.** Number of genome-mapped true peaks at different enrichment score values. **D.** Distribution of genome-mapped true peaks at different enrichment score values. **E.** Number of gene-mapped true peaks at different peak length (bp) values. **F.** Distribution of gene-mapped true peaks at different peak length (bp) values. **G.** Number of gene-mapped true peaks at different enrichment score values. **H.** Distribution of gene-mapped true peaks at different enrichment score values.

### IV. Functional Annotation of Peaks’ Related Gene

The peaks’ related genes are candidate *Ppp1r1b-lncRNA* binding sites, from which we may infer its potential biological impact and mechanisms of function. The true peaks’ related genes, which define *Ppp1r1b-lncRNA*-binding sites, were listed for functional annotation to characterize the functional properties of these genes and their products using GO analysis. At a global level, *Ppp1r1b-lncRNA* binding sites were enriched in molecular function in terms of binding, catalytic activity, transcription regulation, and signal transduction. The nucleus, cellular organelles, and cellular membrane were enriched cellular components. Biological regulation, cellular biogenesis, metabolic process, and cellular transport were enriched biological processes [**Figure 5. A**].

**Figure 5.**
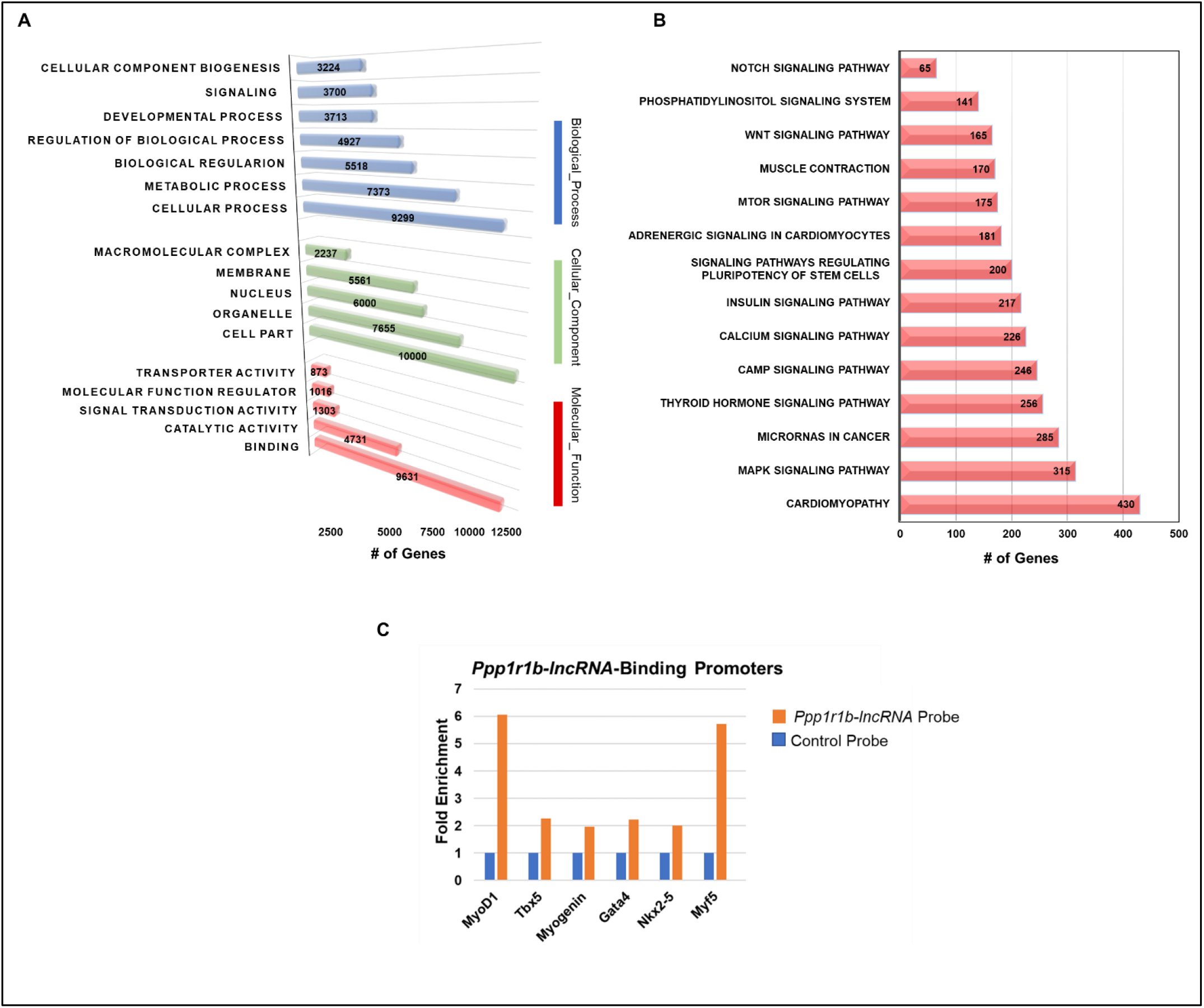
Functional Annotation of *Ppp1r1b-lncRNA*-Binding Sites. **A.** Gene Ontology (GO) analysis of *Ppp1r1b-lncRNA*-ChIRP peaks’ related genes. The top significantly enriched GO terms (FDR < 0.05) involved in biological processes, cellular components, or molecular functions are presented. The number of genes in each term is shown. **B.** Summary of KEGG Pathway analysis of *Ppp1r1b-lncRNA*-ChIRP peaks’ related genes. The top significantly enriched pathways are presented. The number of genes in each pathway is shown. Only true peaks’ related genes are included in these analyses. **C.** *Ppp1r1b-lncRNA*-ChIRP-PCR validation of *Ppp1r1b-lncRNA* interaction with promoters of myogenic differentiation transcription factors.

To elucidate *Ppp1r1b-lncRNA* interactions within certain biological contexts or signaling pathways, KEGG pathway analysis was also performed on all *Ppp1r1b-ncRNA* binding-sites’ related genes [**Figure 5. B**] and [**Table 4**]. Pathways of cardiomyopathy, cancer and pluripotency, transcriptional regulation, and developmental pathways were among the top enriched processes. Further, critical signaling pathways of development and myogenesis, including Wnt, Notch, metabolism, and insulin signaling were among the top enriched signaling pathways.

**Table 4 (Excel Sheet 4):**
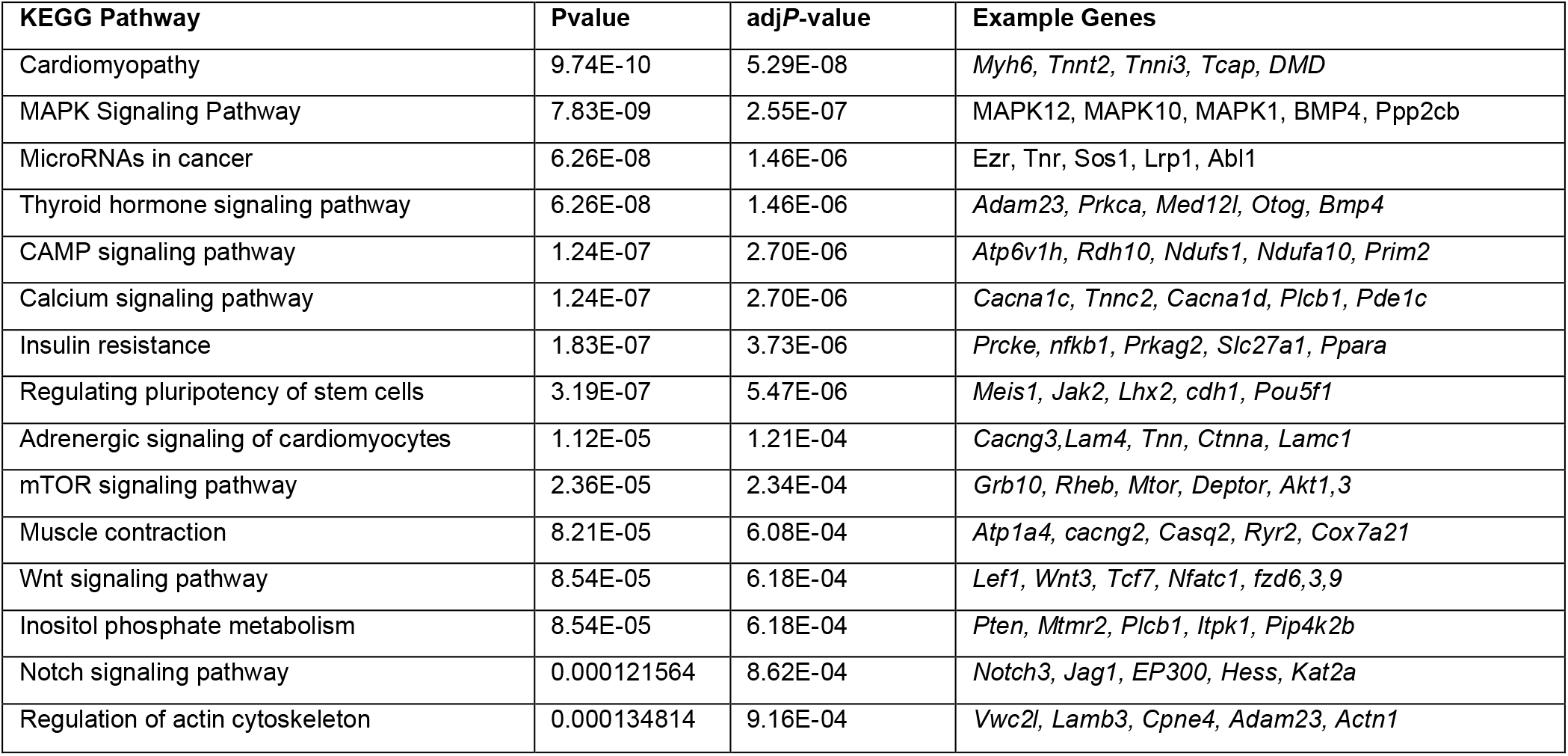
KEGG pathway analysis. Top 20 significant pathways enriched in the peaks’ related genes. *P*-values, adjusted *P*-values, and 5 representative genes in each category are presented.

### V. Promoter Mapped Peaks

As previously demonstrated, *Ppp1r1b-lncRNA* executes its functions through the interaction with promoters of myogenic transcription factors. Further, *Ppp1r1b-lncRNA-*ChIRP peaks (binding sites) are mostly narrow, reminiscent of TF binding sites. Based on these observations we performed an independent ChIRP-PCR. In addition to the previously known interactions with *MyoD1* and *Tbx5*, we validated additional four new interactions with myogenic differentiation factors specific to cardiac and skeletal myocytes identified from ChIRP-seq [**Figure 5. C**]. We further examined *Ppp1r1b-lncRNA* binding to promoter regions using EPDnew [**20, 21**], including all peaks in this analysis. In total, 2871 peaks were mapped to experimentally annotated promoter sequences. Of these,1180 true peaks (enrichment score >=10) were retained as promoter-mapped *Ppp1r1b-lncRNA*-binding sites, accounting for 28% of the true binding sites that mapped to protein-coding genes.

Notably, the promoter-mapped binding sites were significantly enriched with transcription regulators, including those involved in the Wnt signaling pathway (Lef1 and Tcf7), heart muscle development (Gata4 and Mef2c) regulation of transcription by RNA Pol-lI (Sox17, SRF, EGR2) and chromatin modification (Smarcb1 and Taf9). Correspondingly, the binding sites that mapped to these genes had high enrichment scores [**Table 5**]. Intriguingly, up to 80% of the identified promoter-mapped binding sites were enriched with one or more of the previously validated sequence elements of proximal promotes (TATA, transcription initiator, CCAT-box, and GC-box) with established specificity for transcription initiation by RNA Pol-II [**Table 5**]. Together, these results are consistent with *Ppp1r1b-lncRNA* function in transcription initiation of myogenic regulators via binding to their promoter elements.

**Table 5 (Excel Sheet 5):**
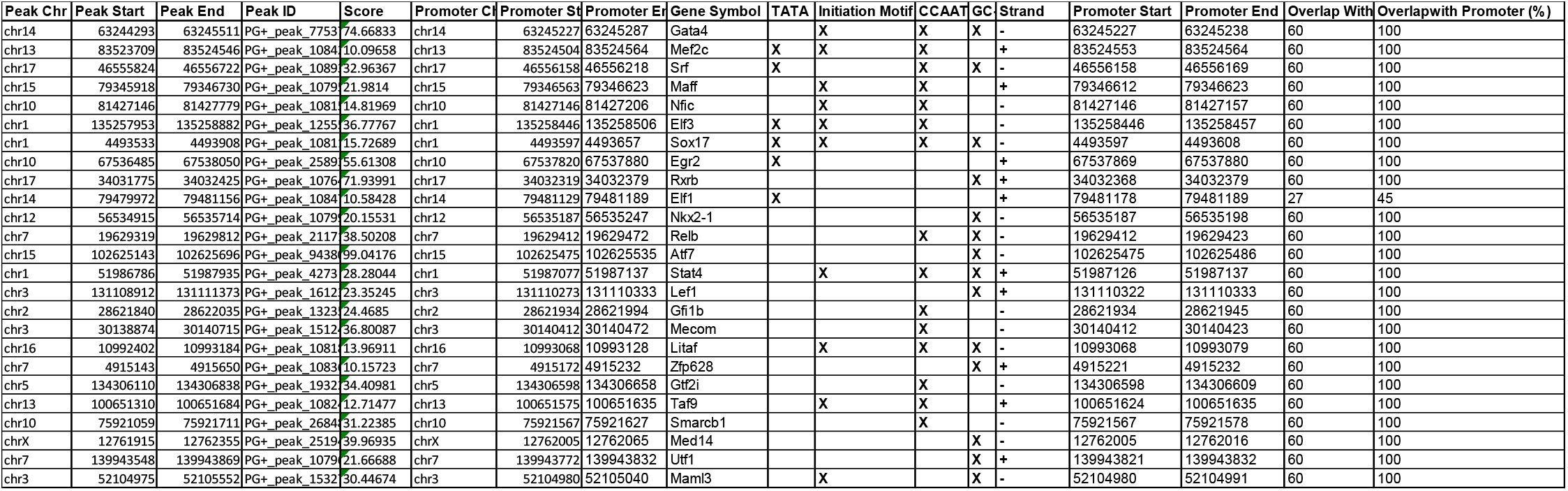
Promoter-mapped *Ppp1r1b-lncRNA*-binding sites involved in transcription by RNA Pol-lI. Top 25 experimentally validated *Ppp1r1b-lncRNA*-bound promoters involved in transaction by RNA Pol-II.

LncRNA binding of transcription factors is mainly governed by their sequence specificity and therefore is typically associated with highly localized ChIRP-Seq signals in the genome. Therefore, we furthered our motif analysis using MEME: Simple Enrichment Analysis, including all *Ppp1r1b-lncRNA*-binding sites. Using this analysis we identified 310 transcription factors with specific motif sequences. Of these transcription factors, 25% belong to the Homeobox family such as HOX and LHX and were enriched with TA-rich motifs such as TTAATTAAT and TAATTA motifs [**Figure 6**] and [**Table 6**]. In addition, a few Zinc finger-related transcription factors were enriched with GC sequence repeats [**Figure 6**] and [**Table 6**]. Together, these findings suggest novel motif sequences for *Ppp1r1b-lncRNA-*specific interactions with transcription factors.

**Figure 6.**
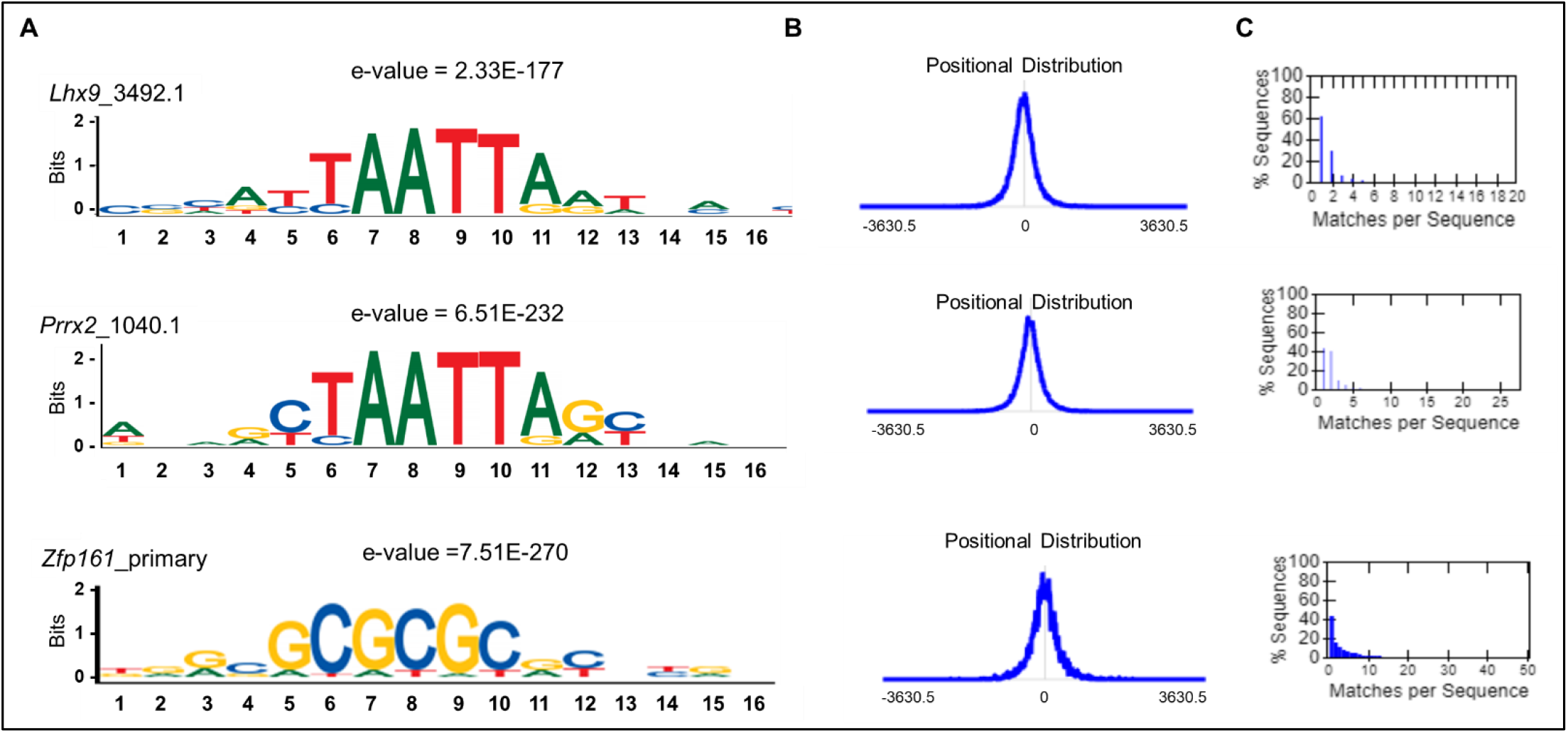
Motif Enrichment in *Ppp1r1b-lncRNA* Binding Sites to Transcription Factors. **A.** Representative examples of significantly enriched motifs in *Ppp1r1b-lncRNA*-binding sites in Homeobox transcription factors (*Lhx9* and *Prrx2* with TA-rich motifs) and zinc fingers (Zfp161 (Zbtb14 with GC-rich motif). **B.** Positional distribution of the best match to the motif in the primary sequences. The plot is smoothed with a triangular function whose width is 5% of the maximum primary sequence length. The position of the dotted vertical line indicates whether the sequences were aligned on their left ends, centers, or right ends, respectively. **C.** The percentage of sequences matching the motif. A sequence is said to match the motif if some position within it has a match score greater than or equal to the optimal score threshold.

**Table 6 (Excl Sheet 6).**
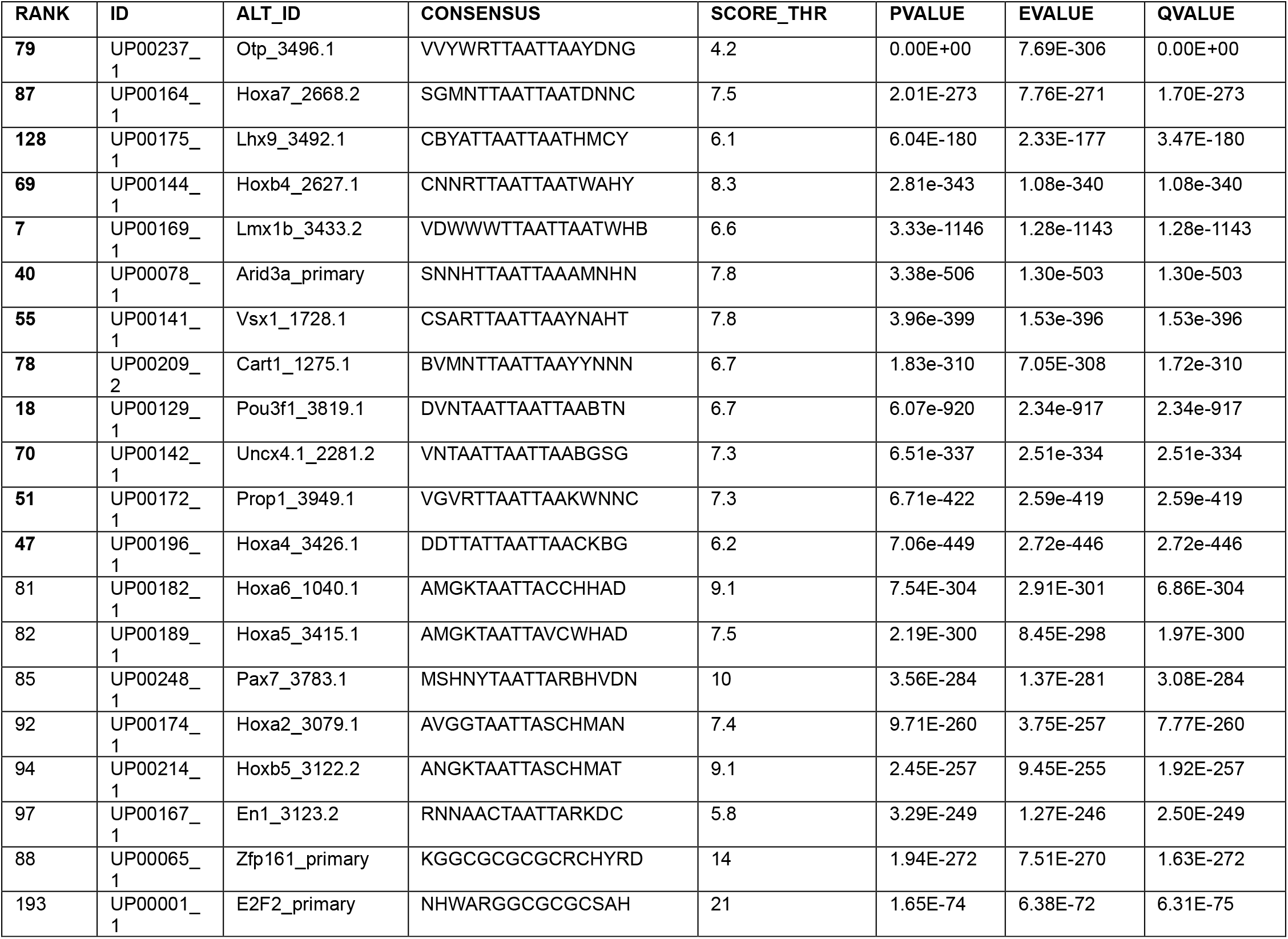
Motif enrichment of Ppp1r1b-lncRNA-bound transcription factors. Top 20 Ppp1r1b-lncRNA-bound transcription factors with enriched motif sequences.

### VI. Enhancer Mapped Peaks

Cell and tissue specificity are governed by tissue/cell-specific enhancer elements. Using Enhancer Atlas 2.0 workflow [**22**], all MACS-derived peaks were mapped to detect the enhancer elements that may be enriched in *Ppp1r1b-lncRNA*-ChIRP signals and to identify their tissue/cell-specific enrichment. In total, more than 12,000,000 enhancer sequences were mapped to all peaks. By applying stringent filtering, only enhancers that are enriched in the true *Ppp1r1b-lncRNA*-binding sites at confidence score ≥1 and ≥30% overlap with a given enhancer were retained, leading to 136521 *Ppp1r1b-lncRNA*-bound enhancer consensuses. Among these signals, 12% (16571 enhancers) showed specific enrichment in cardiac progenitor cells, fetal heart, and limb tissues at high confidence scores [**Table 7**]. These findings correspond to *Ppp1r1b-lncRNA*-specific cellular function in myogenic differentiation of heart and muscle development. Furthermore, histone structure genes and epigenetic modification process were enriched in the enhancers’ enriched *Ppp1r1b-lncRNA*-binding genes.

**Table 7 (Excel Sheet 7).**
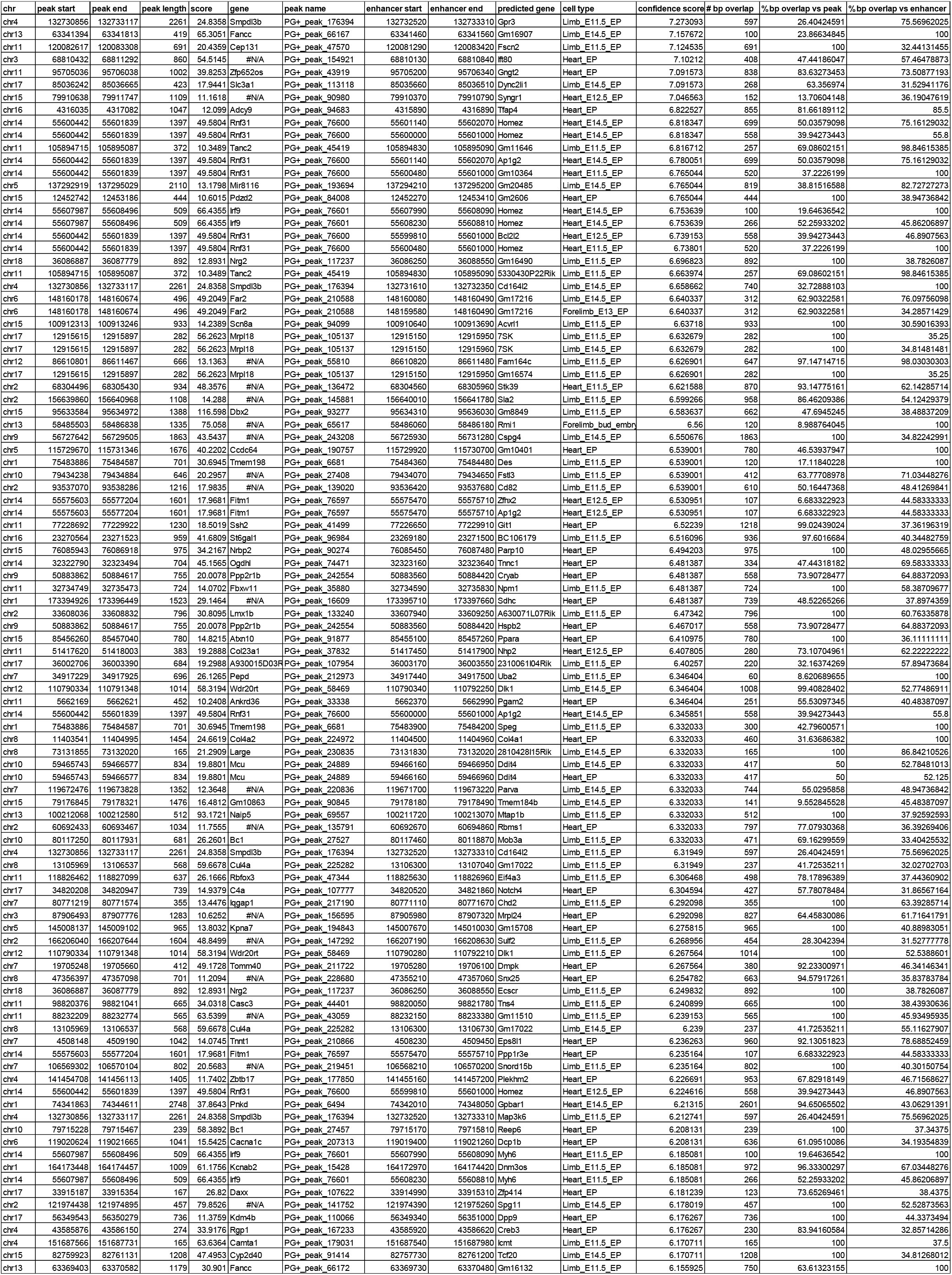
*Ppp1r1b-lncRNA*-bound enhancers. Top 100 Ppp1r1b-lncRNA-enriched enhancers with specificity to fetal heart- or muscle-tissue/cells.

## DISCUSSION

In this study, we applied ChIRP-seq technology against GRCm38/mm10 murine species to identify *Ppp1r1b-lncRNA* chromatin occupancy genome-wide in mouse myoblast cell line, which expresses *Ppp1r1b-lncRNA* [**3,13**]. As described previously [**16,17**], using Glutaraldehyde crosslinking and *Ppp1r1b-lncRNA*-targeted high-affinity probe, the lncRNA-bound DNA sequences were recovered and purified. LacZ probe was used as a negative control, and no cross-hybridization with *Ppp1r1b-lncRNA* was observed. The purified *Ppp1r1b-lncRNA*-ChIRP DNA fragments were used to generate the sequencing libraries and subjected to high throughput single-read sequencing. An Input DNA sample was subjected to the same sequencing protocol and used as a control to allow interpretation of the results.

We selected MACS, a window-based method [**19**], for peak calling based on previous knowledge that *Ppp1r1b-lncRNA* executes its function via the interaction with myogenic transcription factors [**13**]. MACS has been reported to outperform several other methods in the identification of transcription-binding sites that tend to be focal and narrow [**16,19**]. In addition, the MACS pipeline is user friendly and provides important information for each peak, including genomic position, enrichment score, etc.

Using MACS, we identified 244944 Ppp1r1b-lncRNA ChIRP peaks in the genome at *P*<1E-5 and enrichment score >=1. *Ppp1r1b-lncRNA-ChIRP* peaks were focal and narrow averaging 554 bp in length, and typically span less than 300 hundred nucleotides (Mode 165 bp), but occasionally stretched beyond 2K BPs (1% of all peaks). We found the peaks mapped to the intergenic regions (53.6%) and gene regions (47.4%) and distributed on all chromosomes [**Figure 3. C**]. By applying more stringent criteria for detection of true peaks detection, a total of 99732 Ppp1r1b-lncRNA binding sites were detected at high confidence (peak enrichment score >=10), of which 44% mapped to annotated protein-coding genes. Hence, despite applying stringent criteria for true peaks detection, the proportion of binding sites that mapped to the protein-coding genes was retained, and the distribution patterns of the peaks on the different chromosomes remained consistent at different cut-off values for peak length and enrichment scores, both at genes and genome scales.

The enrichments with myogenesis, muscle contractions, and cardiomyopathy in the interacted genes reinforce the essential role of *Ppp1r1b-lncRNA* in myogenic differentiation. Other than myogenic differentiation factors, Wnt signaling, Notch signaling and multipotency pathways are also critical to lineage commitment of skeletal muscle and cardiac progenitors. Moreover, we identified enrichment with sarcomere structures genes (*Myh7*, *Tnnt2*, and *Tcap*) [**Table 4**] and components of myocyte membrane, including the Dystrophin-Glycoprotein complex (DGC) components (*Dmd, Dnta, Dntb*, Sgcd, and *Utrn*), which play important roles in maintaining the integrity of myocyte cellular membranes in heart and skeletal muscles [**Supplemental Table 2**]

The enrichment with ribonucleoproteins and RNA binding proteins (*Hnrnpa1*, *Rbm20*, *Rbfox1*) known to be involved in cardiac and muscle diseases and with chromatin modification genes (*Kdm3b*, *Kdm5c*, and *HDAC4*) supports that *Ppp1r1b-lncRNA* roles may span transcriptional/post-transcriptional regulation and chromatin modifications [**Supplemental Table 2**]. These newly identified candidates at the genomic scale beget further functional studies. In addition, we observed numerous specific *Ppp1r1b-lncRNA*-binding sites that mapped to other annotated regulatory non-coding RNA (*St7*) and micro-RNA (*Mir466* and *Mir1191*) genes of known functions, with signal intensities and enrichment scores comparable to those mapped to the protein-coding genes [**Supplemental Table 2**]. Thus, the current comprehensive repertoire of *Ppp1r1b-lncRNA* occupancy provides a rich resource for a complete understanding of *Ppp1r1b-lncRNA* function.

Importantly, among the *Ppp1r1b-lncRNA-binding* sites, we identified the previously confirmed *Ppp1r1b-lncRNA*-interactions with *TBX5* and *MyoD1* using ChIRP-PCR [**13**] [**Figure 5.C**], supporting that our criteria for detecting *Ppp1r1b-lncRNA*-binding sites can identify true signals with potential functional relevance. Intriguingly, we also detected new unique interactions with other key transcription factors of myogenic differentiation in heart and skeletal muscles and validated these new findings independently using ChIRP-PCR as a gold standard [**Figure 5.C**].

By mapping the *Ppp1r1b-lncRNA* binding sites to the experimentally validated promoters of the EPDnew database we identified 1180 hits located in experimentally validated promoters, within −1000 to +/−200 kb of TSS of a given gene. These signals predict true promoter-mapped *Ppp1r1b-lncRNA-*binding sites based on enrichment score ≥10. Importantly, most of these promoter occupancy sites were enriched with one or more of the four previously annotated regulatory elements that define proximal promoters with binding affinity to RNA Pol-II. This pattern of *Ppp1r1b-lncRNA*-interaction with the promoter’s elements supports the idea that *Ppp1r1b-lncRNA* may promote transcriptional initiation [**29, 30**] [**Table 5**]. This data also demonstrates that ChIRP-seq may precisely uncover biologically relevant interactions.

As stated previously, the observed *Ppp1r1b-lncRNA*-ChIRP peak pattern is like the ChIP-seq peaks of transcription factors binding sites. It also resembles the pattern of HOTAIR-ChIRP, a lncRNA known to recruit PRC2 [**16**]. Like transcription factors, it has been postulated that specific DNA motifs may serve to facilitate lncRNA selective interactions, introducing a new class of regulatory elements in the genome that are specifically targeted by lncRNA. For instance, a GA-rich homopurine motif was previously reported for HOTAIR binding [**16**]. However, unlike HOTAIR, *Ppp1r1b-lncRNA* has been shown to interfere with PRC2 binding at the promoter of myogenic transcription factors [**13**]. Therefore, identifying motifs that infer specificity for *Ppp1r1b-lncRNA* interactions with chromatin may lead, at a mechanistic level, to classify lncRNAs that alter chromatin states in a specific manner (recruiting PRC2 to promoter vs. inhibiting PRC2 binding at promoter). Indeed, using MEME-SEA, we identified TA-rich motifs occupying *Ppp1r1b-lncRNA-*binding sites in the Homeobox family of transcription factors. This finding raises particular interest in future studies, since the Homeobox proteins play critical roles in organogenesis and patterning, including cardiogenesis, during development.

Although *Ppp1r1b-lncRNA-*ChIRP peaks were narrow and focal, they were not restricted to proximal promoters, but a large proportion of them were located within the intronic and intergenic areas, suggesting potential enrichment with other regulatory elements such as enhancers. As distal cis-regulatory elements, enhancers activate the transcription of their target genes in a cell type-specific and tissue-specific manner [**31, 32**]. To date, the EnhancerAtlas 2.0 database [**23**] is the most comprehensive Enhancer database that included 13,494,603 annotated consensus enhancers based on 16,055 datasets in 586 tissue/cell types across nine species. Indeed, we identified significant enrichment with sequence motifs of distal enhancers exceeding 12,000,000. Of these, more than 90% possess cell type/tissue-specificity. By narrowing our analysis to cardiac and limb cell type/tissue, we identified 3390 enhancers at a confidence score > 1 and overlapped with *Ppp1r1b-lncRNA*-binding sites >=30%.

In summary, our study provides *Ppp1r1b-lncRNA* occupancy at a genome-scale. The identified interaction with promoters and enhancers, and their putative enriched motifs may potentially dictate *Ppp1r1b-lncRNA* function in myogenic differentiation and potentially other cellular and biological processes. We should acknowledge the limitations of our study. Despite the comprehensive analysis and the new insights, our study remains descriptive and the biological impacts of the newly identified interactions in altering chromatin state and influencing target gene expression remains to be mechanistically investigated. Pending functional results, selected important candidates would further our understanding of the *Ppp1r1b-lncRNA*-derived functional regulome.

## Data Availability Statement

The datasets used and analyzed during the current study are available from the corresponding author upon reasonable request. The study was registered in the Gene Expression Omnibus (GEO) repository [www.ncbi.nlm.nih.gov/geo] under [Neonatal Heart Maturation (NHM) SupperSeries <u>GSE85728</u>. http://www.ncbi.nlm.nih.gov/geo/query/acc.cgi?acc=GSE85728]. Upon acceptance of this manuscript, ChIRP sequencing data will be deposited to the repository and made publicly available.

## Acknowledgments

We acknowledge the support of the UCLA Children’s Discovery and Innovation Institute, California Center for Rare Disease (CCRD) at the UCLA Institute for Precision Health, the Clinical Genomics Center, and the UCLA Congenital Heart Defects BioCore.

## Authors’ Contributions

Conceptualization, X.K. and M.T.; Methodology, J.H; X.K., and M.T.; software, J.H. and M.T.; validation, X.K.; formal analysis, G.H. and M.T.; investigation, G.H, X.K., C.W., and M.T.; resources, M.T.; data curation, J.H., and M.T.; writing—original draft preparation, J.H. and M.T.; writing—review and editing, J.H., C.W., and M.T.; visualization, J. H., and M.T.; supervision, M.T.; project administration, M.T.; funding acquisition, J.H, and M.T. All authors have read and agreed to the published version of the manuscript.

