## Supplemental File for "Mapping Chromatin Occupancy of *Ppp1r1b-lncRNA* Genome-Wide Using Chromatin Isolation by RNA Purification (ChIRP)-seq"

### TITLE

### AUTHORS AND AFFILIATIONS


John Hwang[1,2,3#], Xuedong Kang[1,2#], Charlotte Wolf[1,4], Marlin Touma[1,2,3,5,6*]

1. Neonatal/Congenital Heart Laboratory, Cardiovascular Research Laboratory, University of California Los Angeles, Los Angeles, CA.
2. Department of Pediatrics, David Geffen School of Medicine, University of California Los Angeles, Los Angeles, CA.
3. Children's Discovery and Innovation Institute, Department of Pediatrics, David Geffen School of Medicine, University of California Los Angeles, Los Angeles, CA.
4. Medical and Life Science, College of Life Science, University of California Los Angeles, Los Angeles, CA.
5. Molecular Biology Institute, College of Life Science, University of California Los Angeles, Los Angeles, CA.
6. Eli and Edythe Broad Stem Cell Research Center, David Geffen School of Medicine, University of California Los Angeles, Los Angeles, CA.

(#) Authors contributed equally to this work.

(*) **CORRESPONDANCE**:
Marlin Touma, MD, PhD.


**Supplemental Table 1. QC results for each sample**

| Sample ID | Clean Reads Number | Clean Reads Q20 Rate (%) | Mapping Rate (%) |
|---|---|---|---|
| Ppp1r1b-lncRNA-ChIRP | 36534935 (pass) | 97.57 (pass) | 95.21 (pass) |
| Control | 38429369 (pass) | 97.67 (pass) | 96.97 (pass) |

Footnotes: Clean Reads Q20 Rate (%): The proportion of bases with quality >= 20 in clean reads. Mappin Rate (%): The proportion of mapping reads in total clean reads.

**Supplemental Table 2. Examples of Genome-wide *Ppp1r1b-lncRNA* Interactions.** Examples of the Dystrophin-Glycoprotein Complex (DGC) genes, Chromatin Modification genes, RNA-binding protein genes, and noncoding RNA genes are presented.

| 1. DGC Complex Componant Genes | | | | | | |
|---|---|---|---|---|---|---|
| **chr** | **peak start** | **peak end** | **peak length** | **score** | **gene** | **peak name** |
| chrX | 84689195 | 84690706 | 1511 | 61.09823 | **Dmd** | PG+_peak_255955 |
| chrX | 84684598 | 84685698 | 1100 | 28.3121 | **Dmd** | PG+_peak_255954 |
| chr18 | 23336549 | 23337457 | 908 | 115.03046 | **Dtna** | PG+_peak_115906 |
| chr18 | 23325157 | 23326210 | 1053 | 84.83178 | **Dtna** | PG+_peak_115905 |
| chr18 | 23392534 | 23393533 | 999 | 41.97489 | **Dtna** | PG+_peak_115909 |
| chr12 | 3717150 | 3717315 | 165 | 93.272 | **Dtnb** | PG+_peak_47995 |
| chr12 | 3718141 | 3718470 | 329 | 83.87779 | **Dtnb** | PG+_peak_47996 |
| chr12 | 3573972 | 3574143 | 171 | 81.42422 | **Dtnb** | PG+_peak_47980 |
| chr10 | 12626678 | 12628059 | 1381 | 79.59871 | **Utrn** | PG+_peak_20393 |
| chr10 | 12854080 | 12854245 | 165 | 70.84226 | **Utrn** | PG+_peak_20427 |
| chr10 | 12694547 | 12695613 | 1066 | 69.15365 | **Utrn** | PG+_peak_20404 |
| chr11 | 47109783 | 47110053 | 270 | 55.0497 | **Sgcd** | PG+_peak_37339 |
| chr11 | 47057251 | 47057626 | 375 | 48.87064 | **Sgcd** | PG+_peak_37330 |
| chr11 | 47048785 | 47049023 | 238 | 48.72734 | **Sgcd** | PG+_peak_37329 |
| chr6 | 4705128 | 4706130 | 1002 | 21.43508 | **Sgce** | PG+_peak_195830 |
| chr6 | 4703533 | 4703834 | 301 | 12.2561 | **Sgce** | PG+_peak_195829 |

| 2. Chromatin Modification Genes | | | | | | |
|---|---|---|---|---|---|---|
| **chr** | **peak start** | **peak end** | **peak length** | **score** | **gene** | **peak name** |
| chr13 | 47073197 | 47073369 | 172 | 76.7658 | **Kdm1b** | PG+_peak_63982 |
| chr13 | 47078702 | 47079862 | 1160 | 73.17467 | **Kdm1b** | PG+_peak_63983 |
| chr13 | 47060585 | 47060984 | 399 | 64.07101 | **Kdm1b** | PG+_peak_63981 |
| chr5 | 122878281 | 122878517 | 236 | 55.13154 | **Kdm2b** | PG+_peak_191770 |
| chr5 | 122980475 | 122981071 | 596 | 26.98227 | **Kdm2b** | PG+_peak_191786 |
| chr5 | 122904528 | 122904693 | 165 | 21.21861 | **Kdm2b** | PG+_peak_191776 |
| chr18 | 34797941 | 34798106 | 165 | 45.58341 | **Kdm3b** | PG+_peak_117053 |
| chr18 | 34807607 | 34808554 | 947 | 39.10465 | **Kdm3b** | PG+_peak_117056 |
| chr18 | 34795128 | 34795663 | 535 | 36.67268 | **Kdm3b** | PG+_peak_117052 |
| chr1 | 134622605 | 134622798 | 193 | 45.85252 | **Kdm5b** | PG+_peak_12446 |
| chr1 | 134631011 | 134632083 | 1072 | 44.7479 | **Kdm5b** | PG+_peak_12447 |
| chr1 | 134615220 | 134615755 | 535 | 17.638 | **Kdm5b** | PG+_peak_12445 |
| chrX | 152237554 | 152237732 | 178 | 22.60131 | **Kdm5c** | PG+_peak_260070 |
| chrX | 152243123 | 152243757 | 634 | 10.4147 | **Kdm5c** | PG+_peak_260071 |
| chr1 | 91982343 | 91982611 | 268 | 38.63383 | **Hdac4** | PG+_peak_8677 |
| chr1 | 92124396 | 92125637 | 1241 | 28.82958 | **Hdac4** | PG+_peak_8701 |
| chrX | 102498812 | 102499094 | 282 | 63.61024 | **Hdac8** | PG+_peak_257267 |
| chrX | 102488025 | 102488534 | 509 | 61.8765 | **Hdac8** | PG+_peak_257266 |
| chrX | 102344693 | 102345122 | 429 | 53.65665 | **Hdac8** | PG+_peak_257251 |
| chr12 | 34280246 | 34280412 | 166 | 67.94064 | **Hdac9** | PG+_peak_50857 |
| chr12 | 34597985 | 34598425 | 440 | 48.45812 | **Hdac9** | PG+_peak_50887 |
| chr12 | 34615678 | 34616011 | 333 | 40.74576 | **Hdac9** | PG+_peak_50888 |
| chr9 | 44808642 | 44808878 | 236 | 30.67642 | **Kmt2a** | PG+_peak_241735 |
| chr9 | 44805749 | 44806212 | 463 | 26.74037 | **Kmt2a** | PG+_peak_241734 |
| chr9 | 44809595 | 44810388 | 793 | 21.4236 | **Kmt2a** | PG+_peak_241736 |
| chr5 | 25392613 | 25393425 | 812 | 50.26575 | **Kmt2c** | PG+_peak_181619 |
| chr5 | 25378141 | 25379054 | 913 | 46.262 | **Kmt2c** | PG+_peak_181618 |
| chr5 | 25276134 | 25276860 | 726 | 39.75114 | **Kmt2c** | PG+_peak_181610 |
| chr15 | 98859225 | 98859745 | 520 | 55.24304 | **Kmt2d** | PG+_peak_93752 |
| chr15 | 98850830 | 98852629 | 1799 | 28.50097 | **Kmt2d** | PG+_peak_93751 |
| chr15 | 98865472 | 98865667 | 195 | 24.1105 | **Kmt2d** | PG+_peak_93753 |

| 3. RNA-Binding Protein Genes |  |  |  |  |  |  |
|---|---|---|---|---|---|---|
| **chr** | **peak start** | **peak end** | **peak length** | **score** | **gene** | **peak name** |
| chr19 | 53855407 | 53855598 | 191 | 139.58006 | **Rbm20** | PG+_peak_128575 |
| chr19 | 53761155 | 53762034 | 879 | 124.10281 | **Rbm20** | PG+_peak_128561 |
| chr19 | 53858364 | 53858531 | 167 | 116.5565 | **Rbm20** | PG+_peak_128576 |
| chr5 | 120195050 | 120195444 | 394 | 70.40952 | **Rbm19** | PG+_peak_191383 |
| chr5 | 120193508 | 120193883 | 375 | 64.84457 | **Rbm19** | PG+_peak_191382 |
| chr5 | 120189785 | 120190481 | 696 | 54.85414 | **Rbm19** | PG+_peak_191381 |
| chr16 | 6414888 | 6416198 | 1310 | 115.828 | **Rbfox1** | PG+_peak_94971 |
| chr16 | 6797396 | 6797702 | 306 | 109.79128 | **Rbfox1** | PG+_peak_95011 |
| chr16 | 6427517 | 6427729 | 212 | 100.58395 | **Rbfox1** | PG+_peak_94972 |
| chr15 | 77233716 | 77234156 | 440 | 74.5144 | **Rbfox2** | PG+_peak_90482 |
| chr15 | 77231644 | 77232467 | 823 | 27.21862 | **Rbfox2** | PG+_peak_90481 |
| chr4 | 154318275 | 154318475 | 200 | 102.92594 | **Prdm16** | PG+_peak_179473 |
| chr4 | 154334323 | 154334857 | 534 | 86.64391 | **Prdm16** | PG+_peak_179474 |
| chr4 | 154452501 | 154455535 | 3034 | 49.83498 | **Prdm16** | PG+_peak_179496 |
| chr15 | 103240706 | 103240977 | 271 | 16.73901 | **Hnrnpa1** | PG+_peak_94488 |
| chr6 | 51467094 | 51467571 | 477 | 44.11124 | **Hnrnpa2b1** | PG+_peak_200295 |
| chr2 | 75662202 | 75663273 | 1071 | 40.07079 | **Hnrnpa3** | PG+_peak_137387 |
| chr14 | 52076109 | 52076847 | 738 | 11.03984 | **Hnrnpc** | PG+_peak_76258 |
| chr6 | 117924842 | 117925050 | 208 | 28.21596 | **Hnrnpf** | PG+_peak_207160 |
| chr6 | 117910485 | 117911022 | 537 | 25.87322 | **Hnrnpf** | PG+_peak_207159 |
| chr17 | 33655634 | 33655904 | 270 | 57.76182 | **Hnrnpm** | PG+_peak_107592 |
| chr17 | 33667655 | 33668158 | 503 | 56.12875 | **Hnrnpm** | PG+_peak_107593 |
| chr19 | 8825251 | 8825865 | 614 | 15.08025 | **Hnrnpul2** | PG+_peak_123617 |

| 4. Noncoding RNA Genes |  |  |  |  |  |  |
|---|---|---|---|---|---|---|
| **chr** | **peak start** | **peak end** | **peak length** | **score** | **gene** | **peak name** |
| chr5 | 76287564 | 76288663 | 1099 | 58.51059 | **Mir1191** | PG+_peak_186787 |
| chr5 | 76354534 | 76354941 | 407 | 38.35189 | **Mir1191** | PG+_peak_186796 |
| chr5 | 76311696 | 76311884 | 188 | 35.35888 | **Mir1191** | PG+_peak_186790 |
| chr1 | 185877241 | 185877408 | 167 | 68.62972 | **Mir297c** | PG+_peak_18130 |
| chr1 | 185722110 | 185722549 | 439 | 67.85114 | **Mir297c** | PG+_peak_18108 |
| chr1 | 185939167 | 185939499 | 332 | 65.92393 | **Mir297c** | PG+_peak_18143 |
| chr2 | 131368673 | 131369289 | 616 | 81.61467 | **Mir3098** | PG+_peak_142936 |
| chr2 | 131295481 | 131295855 | 374 | 71.87949 | **Mir3098** | PG+_peak_142928 |
| chr2 | 131497530 | 131497954 | 424 | 68.29517 | **Mir3098** | PG+_peak_142957 |
| chr14 | 112048326 | 112048720 | 394 | 53.43372 | **Mir466** | PG+_peak_81829 |
| chr14 | 112046628 | 112046887 | 259 | 44.10944 | **Mir466** | PG+_peak_81828 |
| chr14 | 112030977 | 112031235 | 258 | 42.53331 | **Mir466** | PG+_peak_81827 |
| chr11 | 86989885 | 86990051 | 166 | 124.29755 | **Mir8115** | PG+_peak_42897 |
| chr11 | 87069867 | 87070456 | 589 | 110.12631 | **Mir8115** | PG+_peak_42910 |
| chr11 | 86983746 | 86984459 | 713 | 105.99644 | **Mir8115** | PG+_peak_42896 |
| chr6 | 17716033 | 17717095 | 1062 | 65.17329 | **St7** | PG+_peak_196865 |
| chr6 | 17877222 | 17877532 | 310 | 49.70775 | **St7** | PG+_peak_196879 |
| chr6 | 17827628 | 17827793 | 165 | 44.10442 | **St7** | PG+_peak_196877 |
| chr4 | 63619226 | 63619493 | 267 | 105.17794 | **1700018C11Rik** | PG+_peak_169458 |
| chr4 | 63606983 | 63608153 | 1170 | 80.53385 | **1700018C11Rik** | PG+_peak_169457 |
| chr6 | 86453667 | 86454460 | 793 | 103.21735 | **C87436** | PG+_peak_203482 |
| chr6 | 86439699 | 86439891 | 192 | 58.43571 | **C87436** | PG+_peak_203481 |
| chr6 | 86457338 | 86458978 | 1640 | 45.65241 | **C87436** | PG+_peak_203483 |

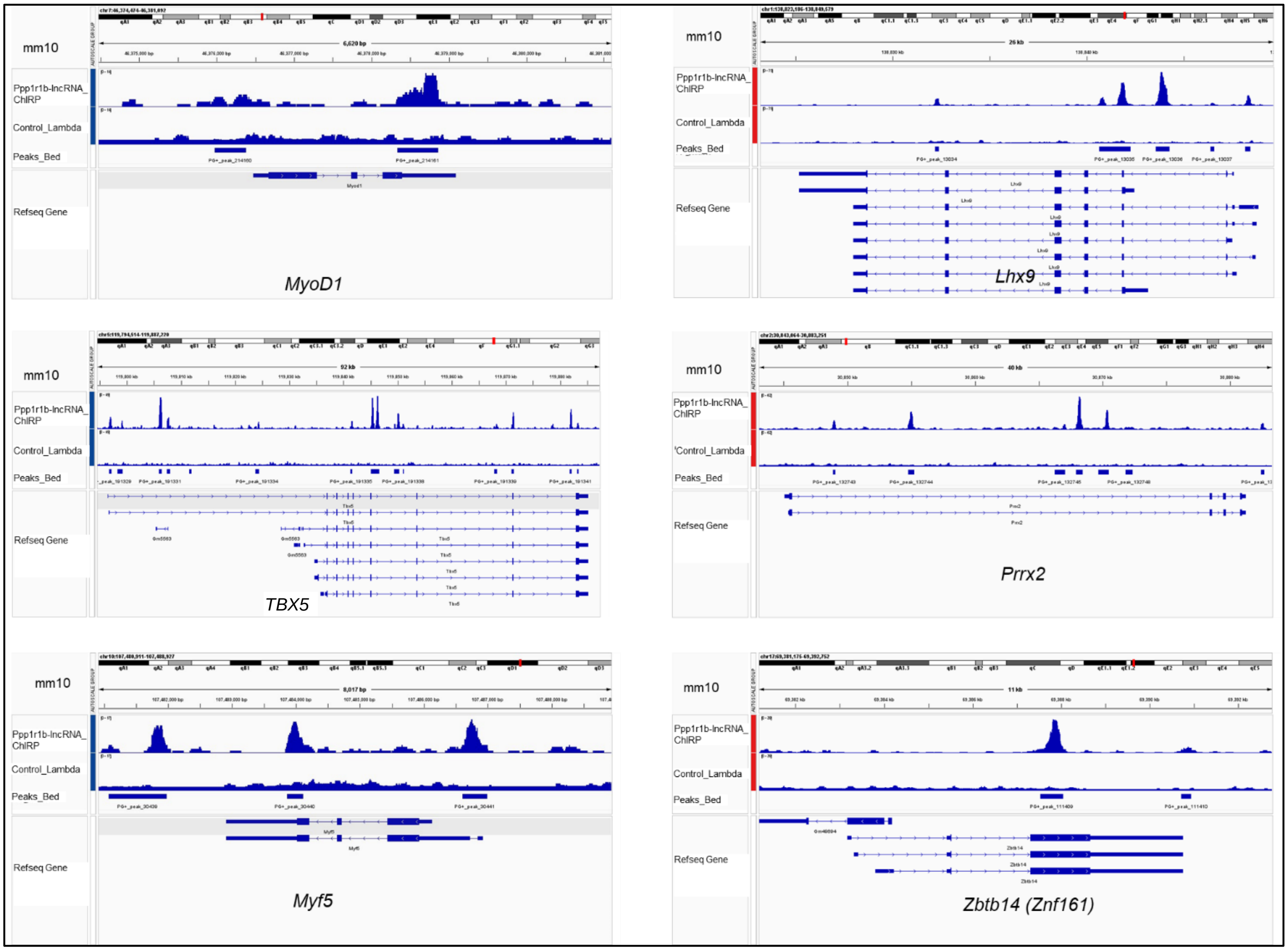


**Supplemental Figure 1. *Ppp1r1b-lncRNA* Binding Sites to Transcription Factors. A.** IGV viewer windows depict *Ppp1r1b-lncRNA*-sites (peaks) enriched in the promoter, intronic, and the distal 50% regions of genes, including myogenic differentiation transcription factors, Homeobox transcription factors (TA-rich motifs) and zinc fingers (GC-rich motif).
